# Dynamics of poly(A) tail length and non-A residues during the human oocyte-to-embryo transition

**DOI:** 10.1101/2021.08.29.458075

**Authors:** Yusheng Liu, Keliang Wu, Fanghong Shao, Hu Nie, Jingye Zhang, Cheng Li, Zhenzhen Hou, Jiaqiang Wang, Bing Zhou, Han Zhao, Falong Lu

**Affiliations:** State Key Laboratory of Molecular Developmental Biology, Institute of Genetics and Developmental Biology, Innovative Academy of Seed Design, Chinese Academy of Sciences, Beijing 100101, China; University of Chinese Academy of Sciences, Beijing 100049, China; Center for Reproductive Medicine, Shandong University, The Key laboratory of Reproductive Endocrinology, Ministry of Education, Shandong University, Jinan 250012, China; State Key Laboratory of Stem Cell and Reproductive Biology, Institute of Zoology, Chinese Academy of Sciences, Beijing 100101, China; Institute for Stem Cell and Regeneration, Chinese Academy of Sciences, Beijing 100101, China; College of Life Science, Northeast Agricultural University, Harbin 150030, China; Department of Life Sciences and Medicine, University of Science and Technology of China, Hefei 230026, China

## Abstract

Poly(A) tail-mediated post-transcriptional regulation of maternal mRNA has been shown to be vital in the oocyte-to-embryo transition (OET) in flies, fish, frogs, and mice^1–8^. However, nothing is known about poly(A) tail dynamics for even a single gene during the human OET, because of the limited availability of human oocytes and embryos in combination with the low sensitivity of previous methods. Here, we systematically profiled the transcriptome-wide mRNA poly(A) tails in human oocytes at the germ-vesicle (GV), metaphase I (MI), and metaphase II (MII) stages, as well as pre-implantation embryos at the 1-cell (1C), 2-cell (2C), 4-cell (4C), 8-cell (8C), morula (MO), and blastocyst (BL) stages using single-oocyte/embryo PAIso-seq1 and PAIso-seq2 methods. We show that poly(A) tail length is highly dynamic during the OET, with BTG4 responsible for global deadenylation. Moreover, we reveal that non-A residues occur primarily in poly(A) tails of maternal RNA, which begin to increase at the MI stage, become highly abundant after fertilization (with U residues occurring in about two thirds, G residues in about one third, and C residues in about one fifth of mRNAs), and decline at the 8C stage. Importantly, we reveal that TUT4/7 can add U residues to deadenylated mRNA, which can then be re-polyadenylated to produce 5′-end and internal U residues. In addition, the re-polyadenylated mRNA can be stabilized through the addition of G residues by TENT4A/B. Finally, we demonstrate that U residues in poly(A) tails mark the maternal transcripts for quicker degradation in 8C human embryos compared to those without U residues. Together, our results not only reveal the dynamics of poly(A) tail length and non-A residues, but also provide mechanistic insights into the regulation of the length and the role of non-A residues during human OET. These findings further scientific understanding and open a new door for studying the human OET.

## Introduction

Poly(A) tails, long chains of adenine nucleotides, are added to the 3′-ends of most eukaryotic mRNAs, where they are essential for mRNA stability and translation^9–16^. The functional importance of poly(A) tail length regulation has been implicated in diverse biological processes and phenomena including gamete development, tumor metastasis, innate immunity, inflammation, obesity, synaptic plasticity, higher cognitive function, and long-term memory^1–3, 16–24^. In addition to regulating poly(A) tail length, non-A residues can be added to the 3′-ends of mRNA poly(A) tails, where they constitute another layer of mRNA turnover regulation^1, 17, 18, 25–31^. Poly(A) tail 3′-end U residues, catalyzed by TUT4/7, promote rapid mRNA decay, whereas 3′-end G residues catalyzed by TENT4A/B can protect mRNA from rapid deadenylation^25, 26^. Interestingly, new technologies have revealed widespread and abundant non-A residues in the internal body of poly(A) tails^27–32^.

The oocyte-to-embryo transition (OET) refers the developmental process from GV oocyte to a matured MII oocyte, followed by fertilization and pre-implantation embryo development in mammals^33–35^. Successful OET is an essential element in mammalian reproduction. Transcription is silent during OET until zygotic gene activation (ZGA). Therefore, all events during the OET are controlled by post-transcriptional regulation of maternal mRNA^33, 35^. For example, global poly(A) tail elongation catalyzed by Wispy during late oogenesis promotes global translation in fly oocytes, while minimal changes in poly(A) tails were observed after egg activation^2–5^. In mammals, only a handful of genes subjected to poly(A) tail regulation have been reported in mice. The functional importance of poly(A) tail-mediated regulation in the OET has been revealed by the fact that deletion of the maternal *Btg4* or *Cnot6l*, which encode core components of the deadenylase complex, in mouse oocytes leads to developmental arrest due to failed deadenylation-mediated degradation of maternal mRNA^36–40^. In addition, *Tut4/7* is required for mouse oogenesis because it facilitates 3′-end U residue-mediated maternal mRNA degradation^18^. Interestingly, zygotes from human female patients with *BTG4* mutations showed a similar phenotype: failed first zygotic cleavage and accumulation of maternal mRNA^41^.

Limitations of poly(A) tail analysis methods have thus far prevented transcriptome-wide poly(A) tail analysis of the mammalian OET. TAIL-seq (or its modified version, mTAIL-seq) and PAL-seq (as well as the associated PAL-seq2) are two technologies based on the Illumina platform that have helped deepen understanding of poly(A) tail length in various organisms^3, 6, 7, 42^. The two main limitations of these methods include the need for microgram-level sample materials (which are impossible to obtain for mammalian oocytes and embryos, particularly humans) and the inability to detect non-A modifications within the main bodies of poly(A) tails (i.e., interior from the 3′ ends). Recently, poly(A) tail analysis using the PacBio platform, including PAIso-seq (called PAIso-seq1 hereafter), PAIso-seq2, and FLAM-seq^27, 28, 32^, has enabled accurate analysis of non-A residues in poly(A) tails, revealing wide-spread U, C, and G residues in the 5′-end, internal, and 3′- end of poly(A) tails, suggesting important regulatory functions of these non-A residues in poly(A) tails. PAIso-seq2 provides the most comprehensive information about poly(A) tails, including accurate tail length as well as the non-A residues, regardless of position, simultaneously^28^. Among these methods, PAIso-seq1 is uniquely sensitive, able to analyze a single mammalian oocyte, while other methods require at least hundreds of input materials for analysis^27^.

The dynamics of the mRNA poly(A) tail, including length and abundance of non-A residues, during the human OET is completely unknown for even a single gene. It is therefore a research priority to explore the poly(A) tail-mediated post-transcriptional regulation during the human OET. Human oocytes and embryos are extremely valuable and limited. PAIso-seq1 is the only method which can provide the necessary sensitivity for comprehensive transcriptome-wide poly(A) tail analysis of these materials, while PAIso- seq2 can be used as a complementary method for validation of the global pattern of poly(A) tails. Here, we applied PAIso-seq1 and PAIso-seq2 to examine the poly(A) tail landscape in human oocytes and early embryos. Our data reveal extensive dynamics of both length and non-A residues of poly(A) tails, as well as the functional and molecular mechanisms underlying these dynamics.

## Results

### Poly(A) tail length dynamics during the human OET

Nothing was yet known about poly(A) tail dynamics during the human OET. The PAIso- seq1 method we recently developed is currently the most sensitive poly(A) tail analysis method, able to analyze single mammalian oocyte, but limited in its ability to analyze RNA with very short or no poly(A) tails at the 3′-ends^27^. The PAIso-seq2 method fills these gaps, with the ability to measure mRNA 3′-ends regardless of their polyadenylation status and to measure non-A residues in poly(A) tails from the 5′-end, internal parts, and 3′-end simultaneously, though it is limited in sensitivity^28^. The PAIso-seq2 method provides fewer informative reads from the same small amount of samples, such as a single oocyte/embryo, than does PAIso-seq1, which is similar to the limitations of the 3′-adaptor ligation-based TAIL-seq method when compared to mTAIL-seq using splint ligation employing A/T base- pairing^3, 6^. PAIso-seq2 is therefore not preferred for extremely low input or single-cell samples. PAIso-seq1, augmented by PAIso-seq2, is thus currently the best available method, with the appropriate sensitivity for transcriptome-wide poly(A) tail analysis of human oocyte and embryo samples.

In order to explore the mRNA poly(A)-tail-mediated post-transcriptional regulation during the human OET, we applied the PAIso-seq1 method to analyze the human oocytes at the germ-vesicle (GV), metaphase I (MI), and metaphase II (MII) stages, as well as pre- implantation embryos at the 1-cell (1C), 2-cell (2C), 4-cell (4C), 8-cell (8C), morula (MO), and blastocyst (BL) stages (Fig.1a), with 3 to 9 single oocytes or embryos for each stage (Extended Data Fig. 1a). Clustering based on the gene expression quantified by PAIso-seq1 showed that the single oocytes or embryos from the same stage clustered together, confirming the quality of the PAIso-seq1 data (Extended Data Fig. 1b).

**Fig. 1.**
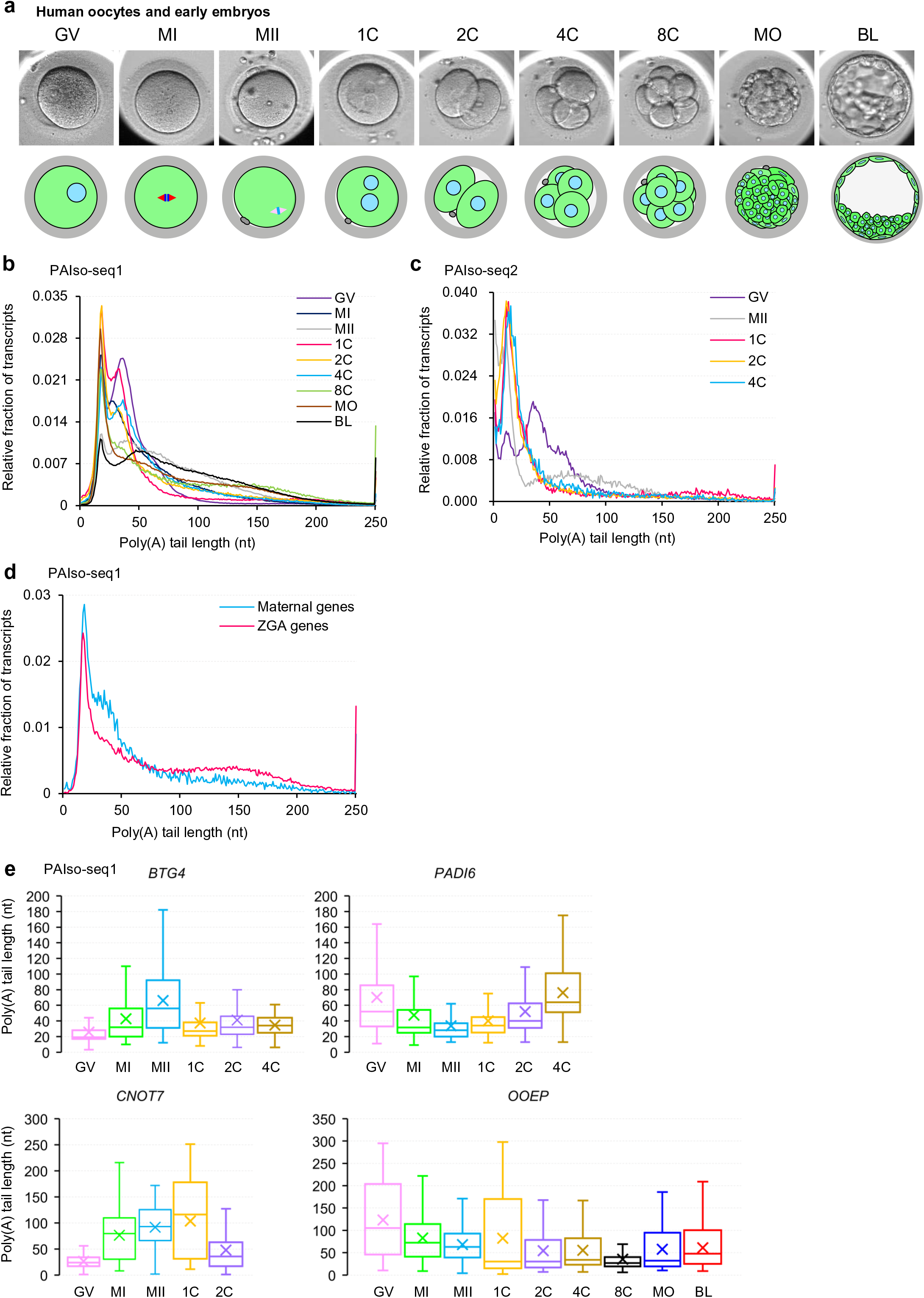
Dynamics of mRNA poly(A) tail length during human OET. **a,** Schematic representation of the process of human oocyte maturation and preimplantation development. GV, germ-vesicle oocyte; MI, metaphase I oocyte; MII, metaphase II oocyte; 1C, 1-cell embryo; 2C, 2-cell embryo; 4C, 4-cell embryo; 8C, 8-cell embryo; MO, morula; BL, blastocyst. **b, c,** Histogram of transcriptome-wide poly(A) tail length in human oocytes and embryos at different stages measured with PAIso-seq1 (**b**) or PAIso-seq2 (**c**). **d.** Histogram of poly(A) tail length of combined transcripts from maternal genes (n = 3,339) or zygotic genes (n = 1,502) in 8-cell stage human embryos measured with PAIso-seq1. **e,** Box plot of the poly(A) tail length of *BTG4*, *PADI6*, *CNOT7*, and *OOEP* in human oocytes and embryos at different stages measured with PAIso-seq1. The “×” indicates mean value, horizontal bars show the median value, top and bottom of the box represent the values of the 25^th^ and 75^th^ percentiles, respectively. Transcripts with poly(A) tails of at least 1 nt for the given gene are included in the analysis. Histograms (bin size = 1 nt) are normalized to cover the same area. Transcripts with poly(A) tail at least 1 nt in length are included in the analysis. Transcripts with poly(A) tail length greater than 250 nt are included in the 250 nt bin.

An overall survey of the data revealed the global poly(A) tail length distribution as well as poly(A) tail length for individual genes are highly dynamic across different stages (Fig. 1b and Extended Data Fig. 3). To further validate the global poly(A) tail dynamics, we also performed PAIso-seq2 for the GV and MII oocytes, as well as 1C, 2C, and 4C embryos (Extended Data Fig. 2). Three to five oocytes or embryos were used for each of the PAIso- seq2 experiments (Extended Data Fig. 2a). The PAIso-seq2 data confirmed the global dynamic changes in poly(A) tail length across stages (Fig. 1c). However, the gene coverage from the PAIso-seq2 data did not allow analysis at the individual gene level due to insufficient unique reads recovered (Extended Data Fig. 4). Therefore, in the current study, the PAIso-seq1 data was used for all analyses, while the PAIso-seq2 data served to validate the key conclusions of the observed global pattern. Both the PAIso-seq1 and PAIso-seq2 data showed that poly(A) tails underwent global deadenylation during human oocyte maturation and experienced global re-polyadenylation after fertilization (Fig. 1b, c), similar to what we observed during the mouse OET^43^. This global dynamic pattern of changes in poly(A) tail length suggests important regulation and function, and this was also the reason we chose the GV, MII, 1C, 2C, and 4C stages for validation by PAIso-seq2.

At the 8C stage, we observed a significant increase in the global poly(A) tail length over that of the 4C stage (Fig. 1b). In human embryos, the major ZGA takes place at the 8C stage^44^. We hypothesized that the newly transcribed mRNA may have longer poly(A) tails, and found that the 8C transcripts from the maternal gene group, which were already much shorter than those of GV oocytes, showed substantially shorter poly(A) tails than those from the ZGA gene group (Fig. 1d).

No studies had yet described gene-level information about poly(A) tails in human oocytes and embryos. In our PAIso-seq1 data, *BTG4* and *CNOT7* showed increases in poly(A) tail length, while *PADI6* and *OOEP* showed decreases in poly(A) tail length during the human oocyte maturation (Fig. 1e). The poly(A) tail length changes associated with these genes are consistent with those described during the mouse oocyte maturation^30, 37, 39^, indicating that these genes are functionally conserved across the OET in both mice and humans, and likely in all mammals.

Because transcription is silent before ZGA during the OET, changes in poly(A) tail length represent differential global post-transcriptional regulation of the maternal transcriptome. Together, the complementary PAIso-seq1 and PAIso-seq2 datasets revealed the first transcriptome-wide dynamic landscape of mRNA poly(A) tail lengths during the human OET, which represents a valuable resource for studying post-transcriptional regulation of human oocytes and pre-implantation embryos.

### Highly dynamic non-A residues in mRNA poly(A) tails during the human OET

Poly(A) tail analysis using the PacBio platform, including PAIso-seq1, PAIso-seq2, and FLAM-seq, enables accurate analysis of non-A residues in poly(A) tails^27, 28, 32^. Interestingly, we found that the proportion of poly(A) tails containing non-A residues is highly dynamic during the human OET, demonstrated across both the PAIso-seq1 and PAIso-seq2 datasets. The global and gene-level proportion of non-A residues increased at the MI stage, with a peak at the 1-cell, 2-cell, and 4-cell stages, and decreased from the 8-cell stage to the blastocyst stage (Fig. 2a, d and Extended Data Figs 5a, 6a). In particular, around two thirds of the poly(A) tails contain U residues at the 1C, 2C, and 4C stages, while G residues are found in about one third, and C residues in only one fifth (Fig. 2a, d). We separated the non-A residues based on their relative positions in poly(A) tails (Fig. 2b). The dynamic changes of non-A residues in both 5′-end and internal parts of poly(A) tails followed a similar pattern across stages in the PAIso-seq1 and PAIso-seq2 data, though PAIso-seq2 is additionally able to detect non-A residues at the 3′-ends (Fig. 2c, e).

**Fig. 2.**
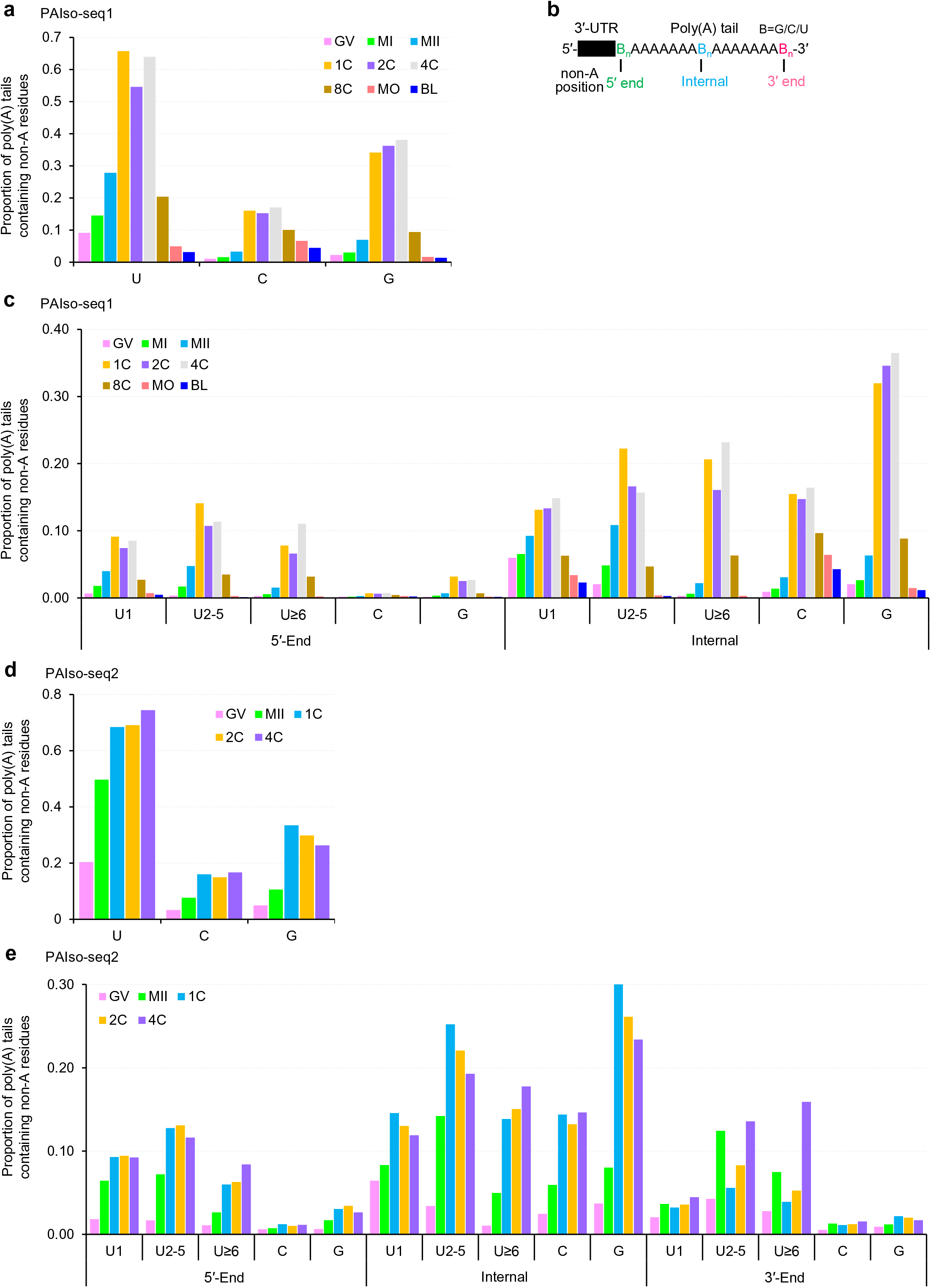
Non-A residues in mRNA poly(A) tails are highly dynamic during human OET. **a, d,** Overall proportion of poly(A) tails containing non-A residues (U, C, or G) in human samples at different stages measured with PAIso-seq1 (**a**) or PAIso-seq2 (**d**). **b**, Diagram of the positions (5′-end, internal, and 3′-end) of non-A residues in poly(A) tails. **c, e,** Proportion of transcripts containing U, C, or G residues at the indicated positions (5′- end, internal, and 3′-end if available) of poly(A) tails in human samples at different stages measured with PAIso-seq1 (**c**) or PAIso-seq2 (**e**). U residues were further divided according to the length of the longest consecutive U sequence (1, 2-5, and ≥6). Transcripts with poly(A) tail at least 1 nt in length are included in the analysis.

U residues are often found in a consecutive manner in poly(A) tails^6, 27^. These sequences of consecutive U residues in poly(A) tails were longer in the human OET than what we observed in the mouse OET^30^. Therefore, we separated the sequences of U residues into three groups (U1, U2-5, U≥6) based on the maximum length of the consecutive U residues.

The three groups showed largely similar dynamics during the human OET in both 5′-end and internal parts of poly(A) tails, with some group specific patterns (Fig. 2c, e). The U1 group showed a pattern similar to that of the G and C residues, increasing at the MI stage, and peaking at the 1C stage, with this high level maintained into the 2C and 4C stages. The U2-5 group began to increase at the MI stage, reaching a peak at the 1C stage, followed by a slight decrease at the 2C and 4C stages. The U≥6 group started to increase at the MI stage and sharply increased at the 1C stage (Fig. 2c, e). Interestingly, deviating from the pattern of U1 and U2-5 groups, the U≥6 group continued to increase in the 4C stage (Fig. 2c). Together with the much more frequent occurrence of U≥6 sequences in poly(A) tails in the human OET than those in the mouse OET, these results suggest distinct regulation of long consecutive U residues (U≥6) in the human OET.

The rapid decline of non-A residues at the 8C stage coincides with ZGA in human embryos, suggesting that the non-A residues are present in the maternal mRNA but not in the newly synthesized zygotic mRNA. Indeed, separating the maternal from the zygotic genes in human 8C embryos revealed that the abundance of non-A residues in maternal genes was much higher than that of zygotic genes (Extended Data Fig. 5b), confirming that the non-A residue dynamics primarily occur in maternal mRNA.

There is no new transcription during the human OET before ZGA; therefore, the observed increase in non-A residues can only be a result of processing existing mRNA poly(A) tails. For a given poly(A) tail containing 5′-end or internal non-A residues, the non-A residues must be first added to the 3′-end, followed by further adenylation to produce a poly(A) tail with 5′-end or internal non-A residues. Interestingly, the pattern of poly(A) tail U residues at the 3′-ends differs from that of the 5′-end and internal parts: it begins to increase at the MI stage and peak at the MII stage, and decreases at the 1-cell stage (Fig. 2c, e). This suggests that massive short or fully deadenylated mRNA 3′ tails appear during oocyte maturation^29^, which can then be uridylated to produce high levels of 3′-end U residues. Next, these 3′-end U residues can be re-polyadenylated to produce 5′-end or internal U residues. Because 5′-end or internal U residues begin to increase at the MII stage, re-polyadenylation must begin at the MII stage and increase greatly after fertilization (Fig. 2c, e).

The U2-5 and U≥6 groups at the 3′-ends showed a second wave of increase in human 4C stage embryos following the decrease observed at the 1C and 2C stages, reaching a level even higher than that of the MII stage (Fig. 2e). This wave of 3′-end U addition tends to generate longer consecutive U (U≥6) tails, which is not seen in the mouse OET^30^. This suggest that the mechanism of this second wave of 3′-end U addition differ from those of the MII stage.

The C and G residues in poly(A) tails begin to increase at the MII stage, and further increase after fertilization, regardless of position at 5′-end, internal parts, or 3′-end (Fig. 2c, e). The abundances of C and G residues in the internal parts are much higher than those in the 5′-end or 3′-end, which are also different from that of U residues (Fig. 2c, e). These suggest that different enzymes and mechanisms control the incorporation of G and C residues versus the U residues. In addition, these differences also suggest different functions of U, C, and G residues in poly(A) tails.

We then measured the length between the end of 3′ untranslated region (UTR) and the longest consecutive sequence of U, C, or G residues in poly(A) tails, which we termed the N (which can be as low as 0) for poly(A) tails with U, C, or G residues (Extended Data Fig. 5c). We observed that the N was short for all types of poly(A) tails with non-A residues (Extended Data Fig. 5d and 6b). The short N (with high levels of tails with 0 N value) suggested that the non-A residues are added to short or fully deadenylated tails. This is consistent with our observation that most re-polyadenylation events occur on partially degraded mRNA 3′-ends^29^. Several poly(A) tail examples with non-A residues in the human 1C PAIso-seq1 samples all showed short N (Extended Data Fig. 5e).

Together, both the PAIso-seq1 and the PAIso-seq2 data reveal that the non-A residues are highly dynamic during the human OET, especially U residues, which are present in the majority of the mRNA transcripts in 1C, 2C, and 4C samples.

### Inhibition of deadenylation results in longer N in human zygotes

We examined the role of global mRNA deadenylation in regulating the length of N in poly(A) tails with non-A residues. We hypothesized that if deadenylation was impaired, this would result in longer N for poly(A) tails with non-A residues, because the substrate for adding non-A residues and re-polyadenylation would be increased with longer poly(A) tails. Btg4 is a critical adaptor protein required to promote global maternal mRNA deadenylation for mRNA decay during mouse oocyte maturation, which is critical for zygotic cleavage after fertilization^37, 39^. Similarly, embryos from human female patients with a *BTG4* mutation also failed in zygotic cleavage after fertilization with accumulation of maternal mRNA^41^. We performed siRNA-mediated knockdown (KD) for candidate factors, including *BTG4*, *TUT4/7*, and *TENT4A/B*, involved in poly(A) tail regulation in GV oocytes, and then matured these oocytes *in vitro* and collected zygotes for PAIso-seq1 analysis after *in vitro* fertilization (Extended Data Fig. 7). The siRNA-mediated KD was successful, as we observed depletion of mRNA of *BTG4*, *TENT4A*, and *TENT4B* in the KD PAIso-seq1 datasets (Extended Data Fig. 7e-f). Expression levels of *TUT4* and *TUT7* were too low to be detected in the siNC sample, which is consistent with the low expression levels of *TUT4* and *TUT7* in the 1C PAIso-seq1 samples, which are sequenced much more deeply (Extended Data Fig. 7b).

The poly(A) tails were longer in the siBTG4 sample compared to other siRNA injected samples (Fig. 3a), confirming the role of BTG4 in global deadenylation. Interestingly, the N for poly(A) tails with U, C, and G residues was longer in siBTG4 but not in other KD samples (Fig. 3b). This is consistent with our observation of reductions in re-polyadenylated degradation intermediates after *BTG4* KD^29^. Therefore, these data confirm that the N for the poly(A) tails with 5′-end and internal non-A residues is regulated by Btg4- mediated global deadenylation, which provides the substrate for the addition of non-A residues.

**Fig. 3.**
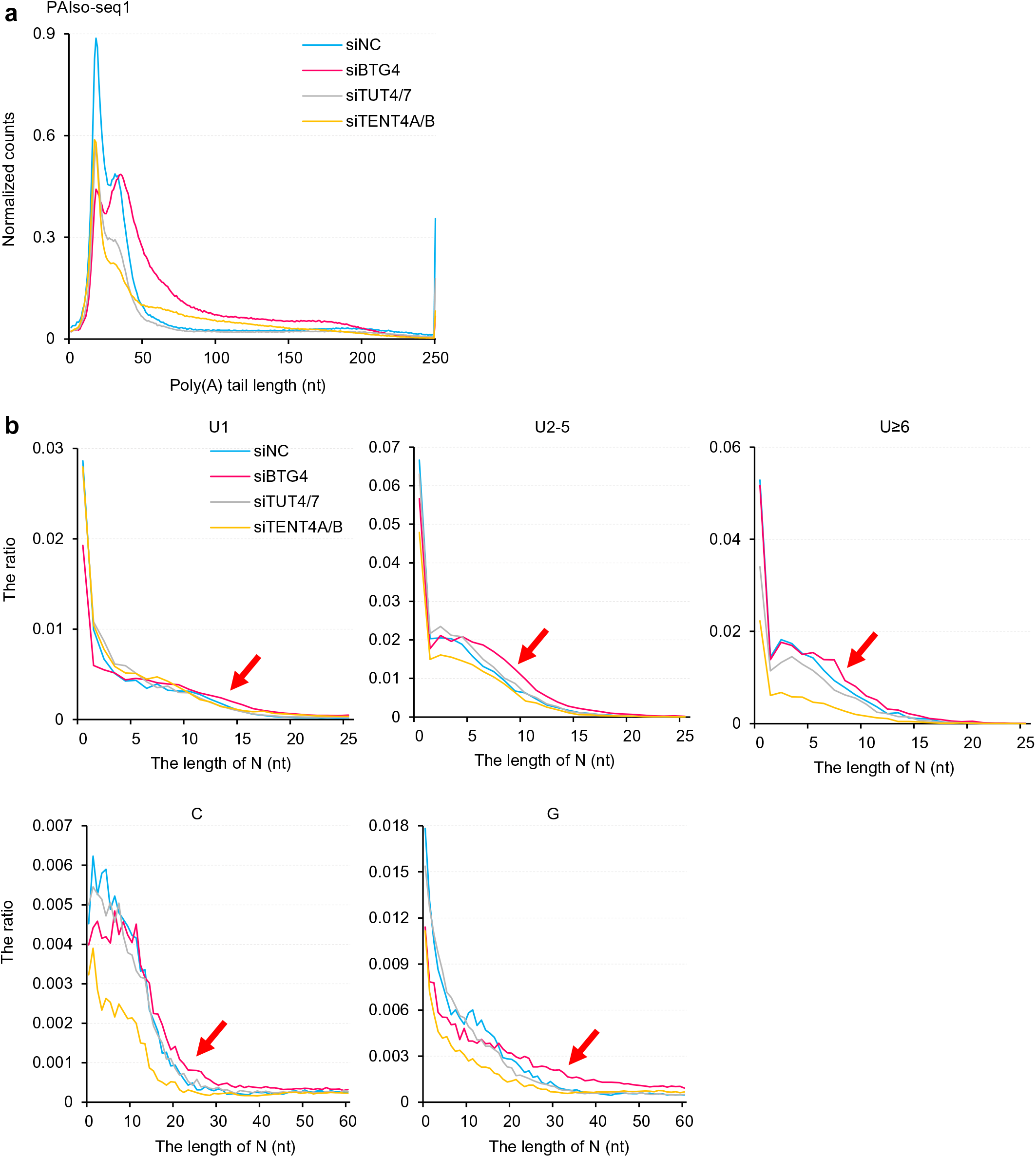
Inhibition of deadenylation results in longer N in human zygotes. **a,** Histogram of poly(A) tail length of all transcripts in *siNC*, si*BTG4*, *siTUT4/7*, or *siTENT4A/B* knockdown (KD) human zygotes measured with PAIso-seq1. Histograms (bin size = 1 nt) are normalized by counts of reads mapped to protein-coding genes in the mitochondria genome. Transcripts with poly(A) tail at least 1 nt in length are included in the analysis. Transcripts with poly(A) tail length greater than 250 nt are included in the 250 nt bin. **b,** Histogram of the length of N (see illustration in Extended Data Fig. 5c) and the ratio of U1, U2-5, U≥6, C, and G residues in *siNC*, si*BTG4*, *siTUT4/7*, or *siTENT4A/B* KD human zygotes measured with PAIso-seq1. Histograms (bin size = 1 nt) are normalized to the total number of transcripts with poly(A) tail at least 1 nt in length. Red arrows highlight the proportion of transcripts with longer N increases upon *BTG4* KD.

### TUT4/7 is responsible for the incorporation of the 5′-end and internal U residues in human zygotes

There is a large number of consecutive U residues at the 5′-end and internal positions (Fig. 2c, e), raising the question of how these are synthesized. TUT4/7 is able to add U residues at the 3′-ends of poly(A) tails^25^. We hypothesized the uridylated 3′-ends can then be further re-polyadenylated to generate 5′-end or internal U residues. To test this hypothesis, we examined the non-A residues in *TUT4/7* KD zygotes. The overall abundance of U residues showed a small decrease in *TUT4/7* KD zygotes (Fig. 4a). We further observed that the decrease in U residues primarily occurred in tails containing 6 or more consecutive U residues (Fig. 4b, red column). This is consistent with the presence of large numbers of long consecutive U residues in human samples. Upon *TUT4/7* KD, uridylation activity at the mRNA 3′-ends decreased, leading to preferential loss of long consecutive U residues. U residues decreased significantly at the individual gene level in TUT4/7 KD zygotes (Fig. 4c). Comparing the number of genes containing different level of U residues, we found that the number of genes with more than 90% of transcripts with U residues decreased the most upon TUT4/7 KD (Fig. 4d). These results reveal that TUT4/7 contributes to the synthesis of 5′-end and internal U residues during the human OET, which is consistent with our observation of Tut4/7 contributing the 5′-end and internal U residues in mice^30^. In humans specifically, there occur more long consecutive U residues which are preferentially affected after *TUT4/7* KD.

**Fig. 4.**
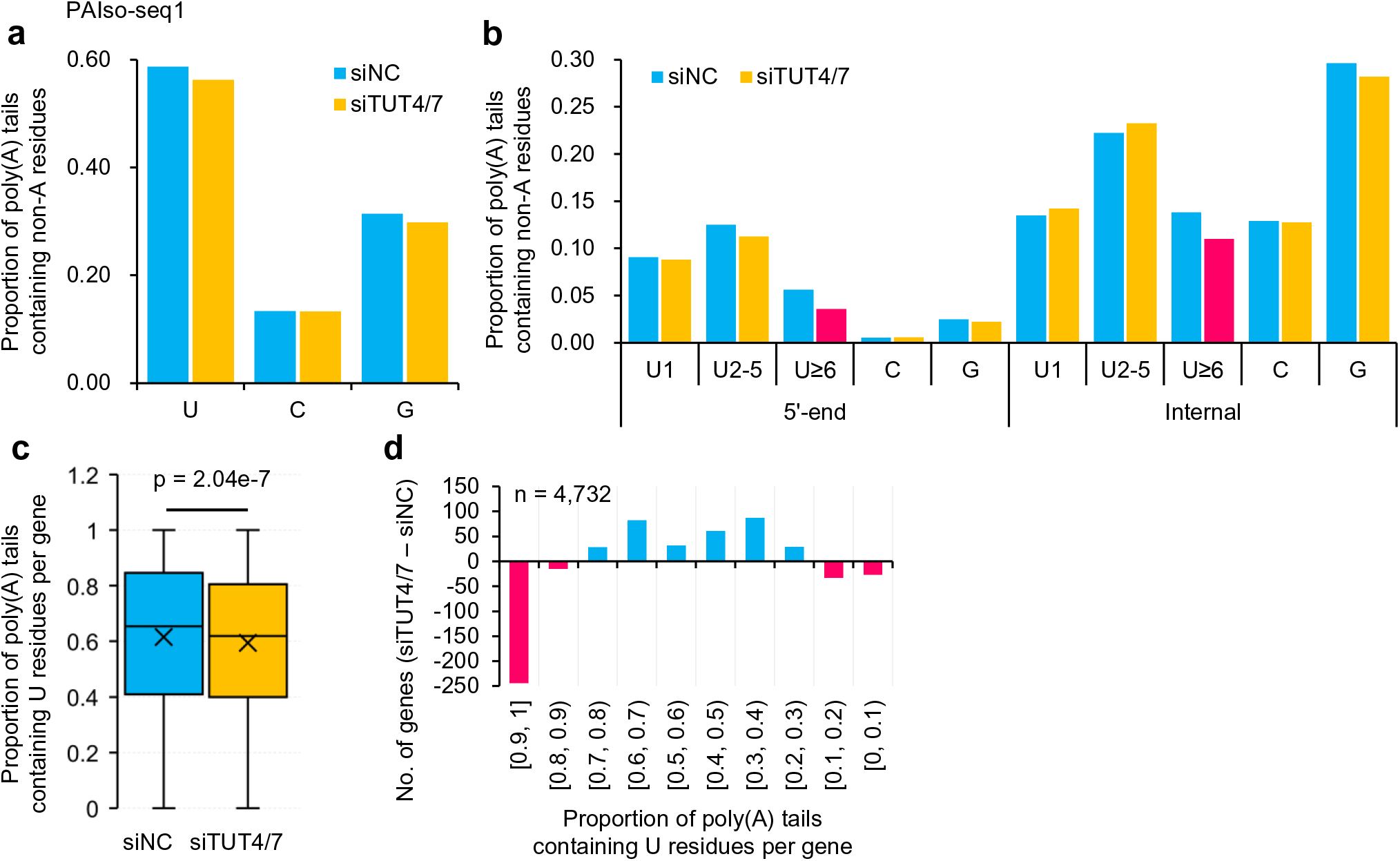
TUT4/7 is responsible for the incorporation of the poly(A) tail 5′-end and internal U residues in human zygotes. **a,** Overall proportion of transcripts containing U, C, or G residues in *siNC* and *siTUT4/7* KD human zygotes measured with PAIso-seq1. **b,** Proportion of transcripts containing 5′-end and internal U, C, or G residues in *siNC* and *siTUT4/7* KD human zygotes measured with PAIso-seq1. U residues were further divided according to the length of the longest consecutive U sequence (1, 2-5, and ≥6). **c,** Box plot of the proportion of reads containing U residues for individual genes (n = 4,732) in *siNC* and *siTUT4/7* KD human zygotes measured with PAIso-seq1. *P* value was calculated with Student’s *t* test. The “×” indicates the mean value, black horizontal bars show the median value, and the top and bottom of the box represent the values of the 25^th^ and 75^th^ percentiles, respectively. **d,** Differences in gene number between *siNC* and *siTUT4/7* KD human zygotes with groups divided by the proportion of reads containing U residues for each gene. Gene number included in the analyses is shown at the top left. Transcripts with poly(A) tail at least 1 nt in length are included in the analysis.

### TENT4A/B protects re-polyadenylated transcripts from degradation by incorporation of G residues in human zygotes

G residues can be incorporated into poly(A) tails by TENT4A/B enzymes in somatic cells, which stabilize the corresponding mRNA by inhibiting deadenylase activity^26^. We observed dynamic incorporation of high level of G residues into poly(A) tails during the human OET (Fig. 2a). The levels of G residues at the 5′-ends and 3′-ends are minimal compared to those of the internal parts of poly(A) tails; therefore, we analyzed G residues as a whole without separating them according to their relative positions. We hypothesized that G residues are catalyzed by TENT4A/B, and can then stabilize the corresponding mRNA. To test this hypothesis, we examined non-A residues in *TENT4A/B* KD human zygotes. The overall levels of G, U, and C residues all decreased in these zygotes (Fig. 5a). Similarly, the levels of G, U, and C residues for individual genes also decreased (Fig. 5b). Moreover, comparing the number of genes containing different levels of G residues, we found that the number of genes with more than 30% of transcripts with G residues decreased most upon *TENT4A/B* KD (Fig. 5c), further confirming the role of TENT4A/B in catalyzing the incorporation of G residues. Next, we examined the consequence of the loss of G residues at the transcript level. The transcript levels of all detected genes showed significant decreases upon *TENT4A/B* KD in human zygotes (Fig. 5d). As a control, KD of *TUT4/7* or *BTG4* in human zygotes did not result in substantial decrease in the levels of G residues or the transcript levels (Fig. 5e-h).

**Fig. 5.**
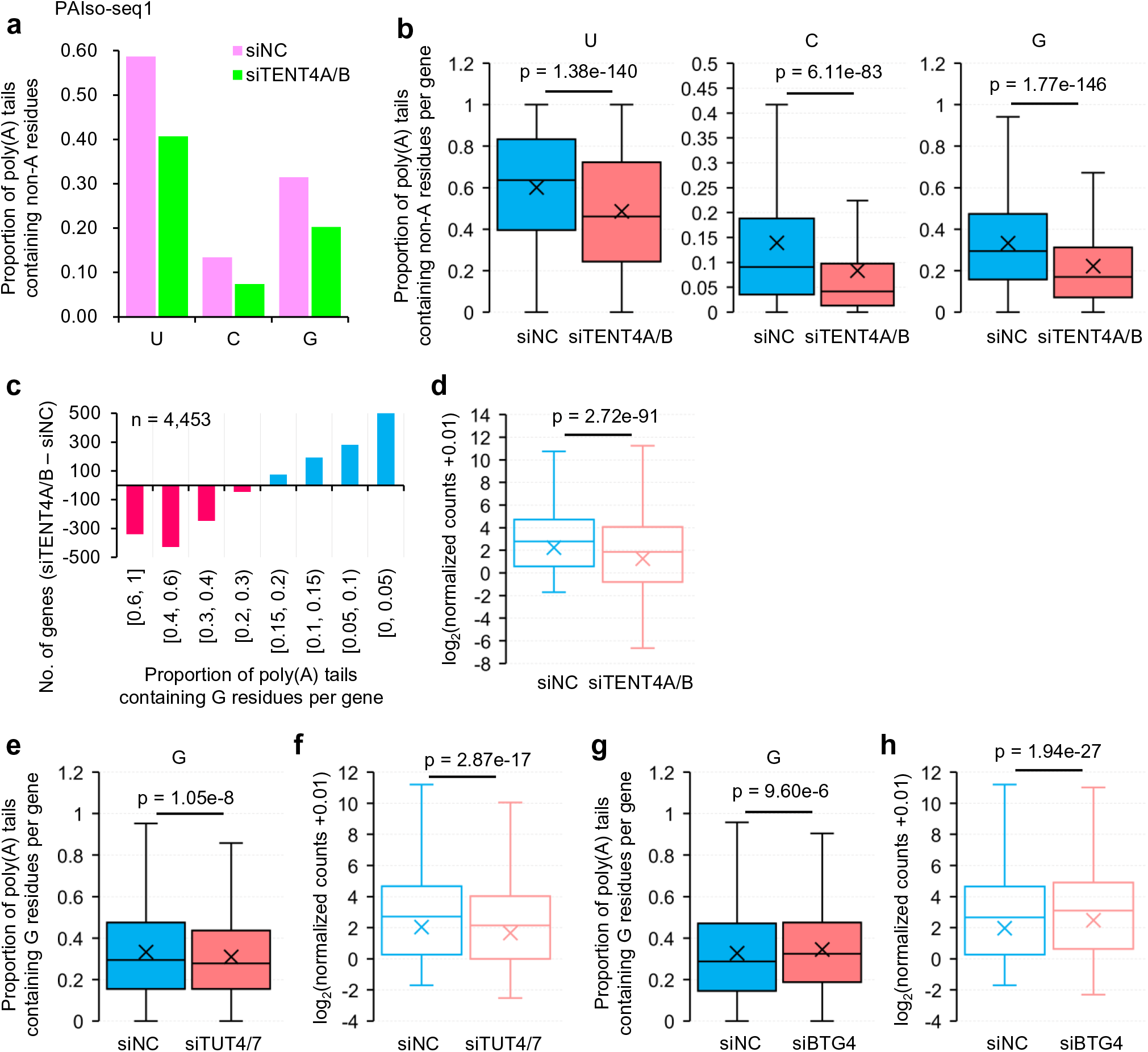
TENT4A/B protects the re-polyadenylated transcripts from degradation by incorporation of G residues in human zygotes. **a,** Overall proportion of transcripts containing U, C, or G residues in *siNC* and *siTENT4A/B* KD human zygotes measured with PAIso-seq1. **b,** Box plot of the proportion of reads containing U, C, or G residues of individual genes (n = 4,474) in *siNC* and *siTENT4A/B* KD human zygotes measured with PAIso-seq1. **c,** Differences in gene number between *siNC* and *siTENT4A/B* human zygotes with groups divided by the proportion of reads containing G residues for each gene. Gene number included in the analyses is shown at the top left. **d, f, h,** Box plots for the normalized counts of genes in *siNC* and *siTENT4A/B* (**d**, n = 11,149), *siNC* and *siTUT4/7* (**f**, n = 11,393), or *siNC* and *siBTG4* (**h**, n = 11,504) KD human zygotes measured with PAIso-seq1. **e, g,** Box plots of the proportion of reads containing G residues of individual genes in *siNC* and *siTUT4/7* (**e**, n = 4,732) or *siNC* and *siBTG4* (**g**, n = 5,035) KD human zygotes measured with PAIso-seq1. All *p* values were calculated with Student’s *t* tests. For all box plots, the “×” indicates the mean value, horizontal bars show the median value, and the top and bottom of the box represent the values of the 25^th^ and 75^th^ percentiles, respectively. Transcripts with poly(A) tail at least 1 nt in length were included in all analyses of proportion of reads containing non-A residues. The read counts were normalized by counts of reads mapped to protein- coding genes in the mitochondria genome if normalization was indicated.

The effects of *TENT4A/B* KD on the G residue levels and transcript levels in human zygotes is stronger than those observed in *Tent4a/b* KD mouse MII oocytes^30^. These differences are most likely due to the different developmental stages analyzed. The level of G residues was much higher at the 1C than at the MII stage in both human and mouse. Therefore, there were many fewer genes with high levels (≥30%) of G residues in the MII sample than in the 1C sample, while genes with high levels of G residues are those preferentially affected by *TENT4A/B* KD in both mouse and human samples. Therefore, a much greater number of genes are affected by *TENT4A/B* KD in human zygotes than in mouse MII oocytes. In addition, the abundance of U and C residues also decreased upon *TENT4A/B* KD in human zygotes, suggesting that the role of TENT4A/B is not specific to poly(A) tails with G residues, but common to all types of re-polyadenylated poly(A) tails. Consistent with the effect of *TENT4A/B* KD on global non-A residues, our unpublished results showed that *Tent4a* KO female mice had reduced fertility.

Together, these results reveal that TENT4A/B contributes to the incorporation of G residues, which stabilize the re-polyadenylated transcripts during the OET in both humans and mice.

### Maternal transcripts with U residues degrade faster than those without U residues

Transcripts with U residues are the dominant polyadenylated mRNA in human 1C, 2C, and 4C embryos, suggesting important functions for these U residues. In mammals, TUT4/7 regulates mRNA degradation during mouse oogenesis^18^. In addition, zygotic KD of *Tut4/7* mRNA leads to defects in maternal mRNA clearance in mouse embryos^41^. However, the role of the U residues added between the GV stage and fertilization is unknown. The 5′- end and internal U residues decrease substantially in 8C human embryos, in which ZGA and global maternal mRNA clearance take place. Therefore, we tested whether the transcripts with or without U residues from maternal genes degraded at a similar rate. Interestingly, the transcript levels of mRNA with or without U residues for each maternal gene revealed that the maternal mRNA with U residues degraded faster than those without U residues from the 4C to the 8C stage (Fig. 6a). We further observed that the ratio of transcripts with U residues to those without U residues decreased significantly for maternal genes from 4C to 8C stage (Fig. 6b), further confirming that mRNA with U residues degraded faster in human 8C embryos.

**Fig. 6.**
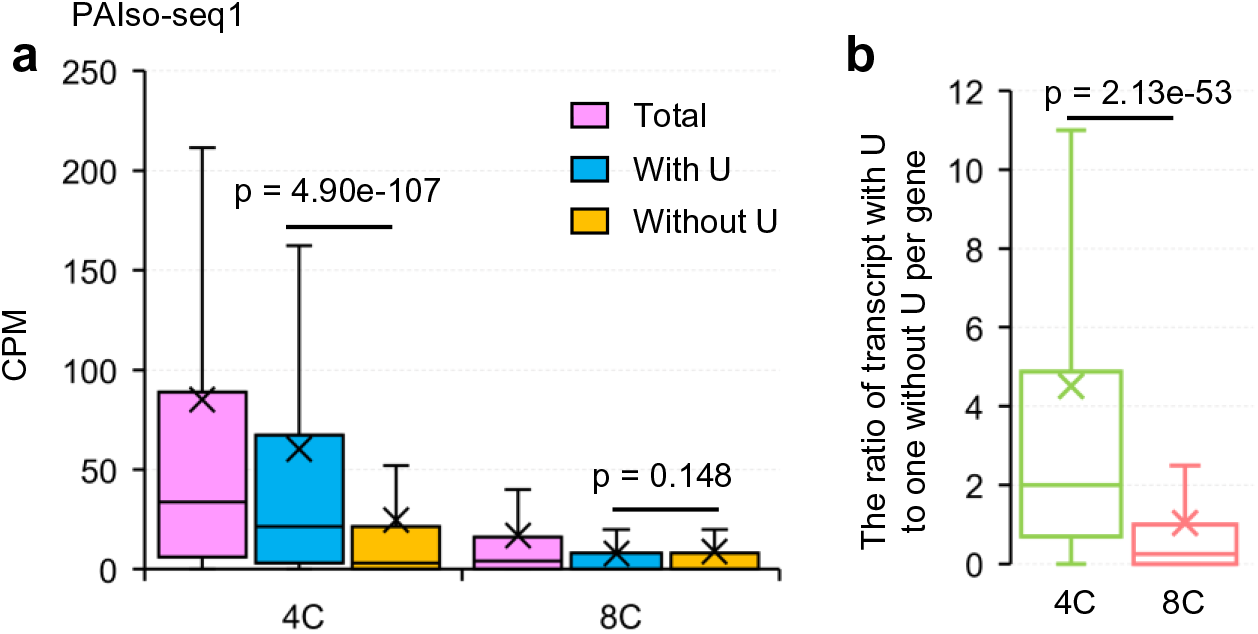
Maternal transcripts with U residues degrade faster than those without U residues in human 8C embryos. **a,** Box plot of the level of total transcripts, transcripts with U residues, and transcripts without U residues for each maternal gene (n = 3,339) in human 4C and 8C embryos measured with PAIso-seq1. Transcripts with poly(A) tail at least 1 nt in length are included in the analysis. **b,** Box plot of the ratio of number of transcripts with U residues to those without U residues for each maternal gene in human 4C and 8C embryos measured with PAIso-seq1. All *p* values were calculated by Student’s *t* tests. For all box plots, the “×” indicates the mean value, horizontal bars show the median value, and the top and bottom of the box represent the values of the 25^th^ and 75^th^ percentiles, respectively.

5′-end and internal U residues promote rapid maternal mRNA degradation at the time of ZGA during the human OET, which is consistent with our findings in mice, rats, and pigs^30, 31^. Although the time between fertilization and ZGA differs substantially between mice, rats, pigs, and humans^44^, the 5′-end and internal U residues serve similar functions during the OET, marking the maternal mRNA and promoting rapid degradation at the stage when ZGA takes place for completion of the OET.

Tut4/7-mediated uridylation has been demonstrated to be essential for mouse reproduction. *Tut4/7* KO during oogenesis and *Tut4/7* KD in the fertilized egg both cause pre-implantation defects in mice^18, 41^. The role of TUT4/7 in human reproduction was completely unknown before this study. Together with the findings in the current study, this shows that U residues added into poly(A) tails by TUT4/7, regardless of their relative positions in poly(A) tails, all ultimately promote mRNA degradation. A previous study revealed that 3′-end U residues are tightly coupled to degradation^25^. Surprisingly, U residue addition is temporally decoupled from immediate mRNA degradation, and can be re-polyadenylated during the OET until degradation, when ZGA takes place in humans, mice, rats, and pigs.

## Discussion

Nothing was yet known about poly(A) tail dynamics during the human OET. In this study, using PAIso-seq1 and PAIso-seq2, we obtained transcriptome-wide poly(A) tail data, including length and non-A residues, from the GV stage oocyte to the blastocyst stage, providing a rich resource for studying post-transcriptional regulation during the human OET (Fig. 7a). In addition, we revealed the critical roles of BTG4, TUT4/7, and TENT4A/B in regulating transcriptome-wide poly(A) tails (Fig. 7b-d). These findings facilitate further understanding of post-transcriptional regulation of the human OET.

**Fig. 7.**
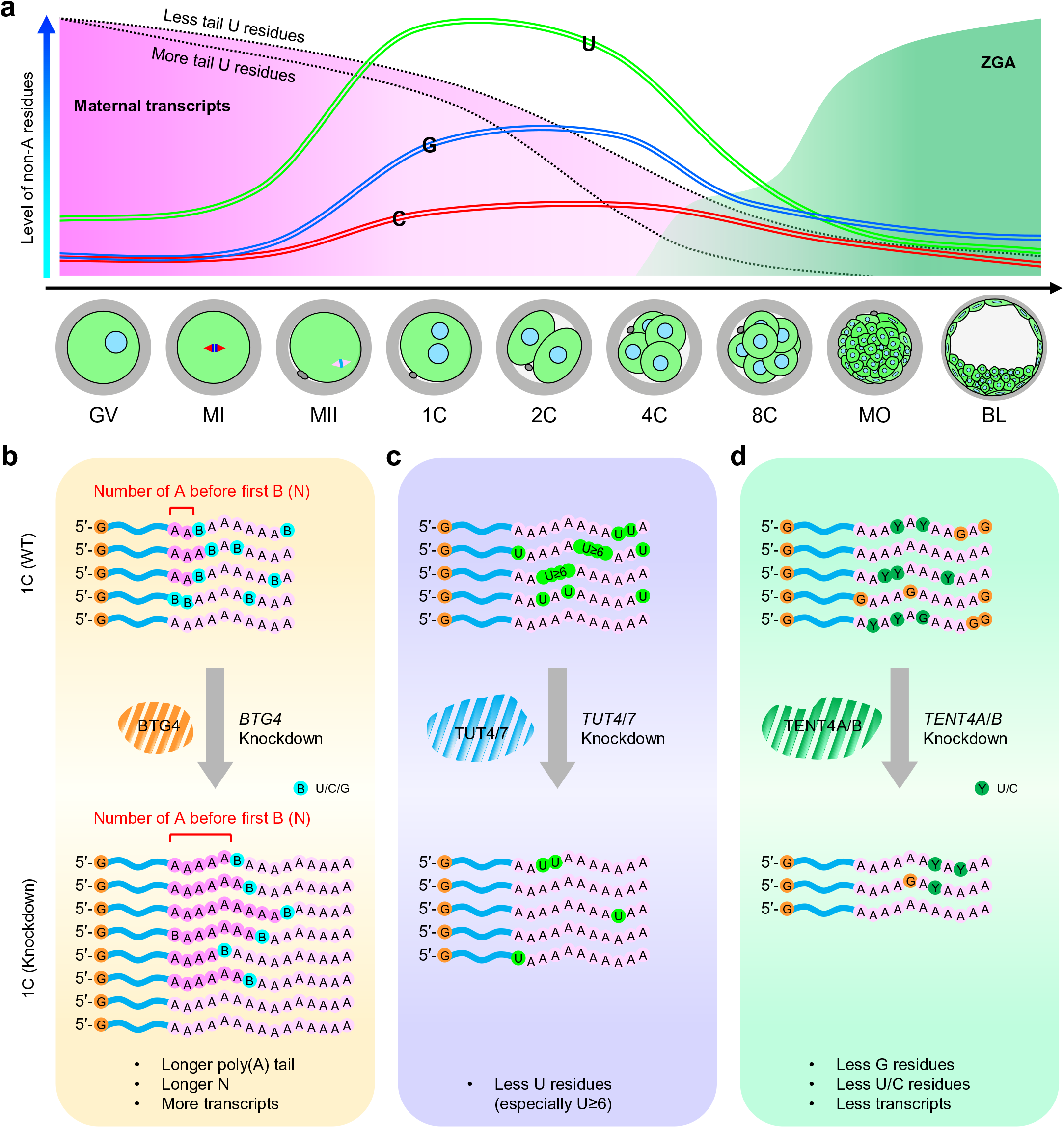
Summary of poly(A) tail dynamics during the human OET. **a,** Illustration of dynamics of maternal transcripts, ZGA transcripts, and non-A (U, C, and G) residues during the human OET. The U residues mark maternal transcripts for faster degradation. **b-d,** Illustration of the effects of *BTG4* (**b**), *TUT4/7* (**c**), or *TENT4A/B* (**d**) knockdown on the metabolism of poly(A) tails and the stability of mRNA transcripts during the human OET.

Poly(A) tail dynamics are integral in development across taxonomic groups. Global poly(A) tail elongation mediated by Wispy during late oogenesis is an essential event for the OET in *Drosophila*, while minimal changes in poly(A) tails were observed after egg activation^2–5^. We revealed that in humans, maternal mRNA typically undergoes shortening of the poly(A) tails in oocyte maturation, while global mRNA re-polyadenylation is seen primarily in zygotes after fertilization. This observation is consistent with our recent observations of the OET in mice^43^. However, deletion of *Tent2*, the mouse homolog of *Wispy*, does not affect re-polyadenylation or embryo development in the mouse OET^45^, suggesting other non-canonical poly(A) polymerases might be responsible for the global re-polyadenylation observed in the mammalian OET. Our unpublished data showed that the fertility decreased in *Tent4a* KO female mice. Therefore, it is likely that Tent4a/b can contribute to both the addition of G residues and the global re-polyadenylation in mammals, which presents an interesting direction for future studies. These disparate observations indicate that global mRNA poly(A) tail length regulation differs between vertebrates and invertebrates.

Maternal mRNA is quickly cleared after ZGA. Interestingly, if maternal RNA deadenylation is blocked by treatment with adenosine monophosphate (AMP) (Extended Data Fig. 8a), which inhibits the general active deadenylation of poly(A) tails by powerfully inhibiting the major de-adenylating enzymes Cnot6l and Cnot7^46^, maternal mRNA with U residues accumulates, while the transcript level of ZGA genes decreases (Extended Data Fig. 8b-d). Therefore, precise regulation of maternal mRNA clearance at the correct timing is critical for ZGA. The U residue-containing poly(A) tails are primarily synthesized in the MII stage, and do not degrade until the 2C stage in mice and rats, the 8C stage in human embryos, and the MO stage in pig embryos, ^30, 31^, indicating a conserved mechanism of stable mRNA with U residue-containing poly(A) tails before ZGA. Transcripts with poly(A) tail 5′-end and internal U residues account about two thirds of the polyadenylated maternal mRNA transcripts in 1C, 2C, and 4C human embryos. These U residues are incorporated by TUT4/7. There is no new transcription at these stages; therefore, these transcripts with U residues are synthesized in the 1C embryo, and are maintained stably until at least the 4C stage, which takes nearly two days. This is surprising, given that previous work showed that TUT4/7-mediated uridylation is known to be coupled with quick mRNA degradation^25^. We found that the transcripts with U residues maintain a high level in the 1C, 2C, and 4C stages. Therefore, our results reveal that the mRNA uridylation at 3′-ends can be temporally decoupled from degradation during the OET. The U residues incorporated by TUT4/7 do not lead to immediate degradation, but serve as marks in transcripts for degradation at a later stage. Identifying the stage-specific biochemical mechanisms responsible for stabilization versus degradation of transcripts with these U residues presents an interesting research direction for future studies.

In this study, we revealed extensive poly(A) tail dynamics and regulation during the human OET. Because poly(A) tails are universal in mRNAs, poly(A) tail length and non-A residue mediated post-transcriptional regulations could be general mechanisms that control diverse biological or disease processes.

## Materials and Methods

### Human oocytes and embryos

The use of human gametes and embryos for this research follows the Human Biomedical Research Ethics Guidelines (set by National Health Commission of the People’s Republic of China on 1 December 2016), the 2016 Guidelines for Stem Cell Research and Clinical Translation (issued by the International Society for Stem Cell Research, ISSCR), and the Human Embryonic Stem Cell Research Ethics Guidelines (set by China National Center for Biotechnology Development on 24 December 2003). All human-related experiments in this study are in compliance with these relevant ethical regulations. The aim and protocols of this study have been reviewed and approved by the Institutional Review Board of Reproductive Medicine, Shandong University.

The donor women were 25–38 years old with tubal-factor infertility and with partners with healthy semen. Written informed consent was obtained from all oocyte and sperm donors. Oocytes were donated from patients receiving *in vitro* fertilization treatments. For early embryos, the donated oocytes were fertilized using donated sperm by intracytoplasmic sperm injection (ICSI). Oocytes and embryos were randomly assigned to experimental groups. All embryos were collected under standard clinical protocols. Single oocytes or embryos were used for PAIso-seq1 analysis with 3 – 9 replicates for each stage (details in Extended Data Fig. 1a). 3 – 5 oocytes or embryos were used for each PAIso- seq2 replicate (details in Extended Data Fig. 2a). All embryos used in this study were cultured no more than 7 days and only used for molecular research analyses.

### Drug treatment of human oocytes and embryos

AMP (Sigma, 2 mM final concentration) and 3′-dA (Sigma, 2 mM final concentration) were dissolved in G1.5 medium (Vitrolife). A medium without drug was used as a control. α-Amanitin was dissolved in DMSO. The final concentration for α-Amanitin treatment was 1 mg/mL. An equal amount of DMSO was added to the medium as the control for the α- Amanitin treatment. For treatment, human zygotes were cultured directly in medium containing the indicated drugs immediately after intracytoplasmic sperm injection and collected at the 8C stage.

### PAIso-seq1 and PAIso-seq2 library construction

The PAIso-seq1 libraries for human single oocytes or embryos were constructed following the single-cell PAIso-seq1 protocol as previously described^27^. PAIso-seq2 libraries were constructed with 3 - 5 human oocytes/embryos following the PAIso-seq2 protocol described in another study with minor modifications^28^. The oocytes/embryos were lysed in buffer containing 0.5 μl of RNase inhibitor and 10.5 μl of 0.2% (vol/vol) Triton X-100 at 80°C for 5 min. The lysate was used directly for the 3′-adaptor ligation following the PAIso-seq2 protocol. The libraries were size selected by Pure PB beads (1x beads for cDNA more than 200 bp and 0.4x beads for cDNA more than 2 kb; the two parts of the sample were combined at equal molarity for further library construction), and made into SMRTbell Template libraries (SMRTbell Template Prep Kit). The libraries were annealed with the sequencing primer and bound to polymerase, and finally the polymerase-bound template was bound to Magbeads and sequenced using PacBio Sequel or Sequel II instruments at Annoroad.

### PAIso-seq1 and PAIso-seq2 sequencing data processing

The PAIso-seq1 and PAIso-seq2 data were pre-processed following our recently published procedures^30^. The human genome annotation used in the analysis was gencode.v36 from Ensembl. The clean CCS reads resulted from the pre-processing procedures were then ready for downstream analysis.

### Poly(A) tail sequence extraction

The poly(A) tail sequence extraction follows our recently published procedures^30^. Clean CCS reads were aligned to human reference genome (GRCh38) using minimap2 (v.217- r941) with the following parameters “-ax splice -uf --secondary=no -t 40 -L --MD --cs -- junc-bed Homo_sapiens.GRCh38.102.gtf.bed”^47^. Alignments with the “SA” (supplementary alignment) tag were ignored. The terminal clipped sequence of the CCS reads in the alignment bam file was used as candidate poly(A) tail sequence. We defined a continuous score based on the transitions between the two adjacent nucleotide residues throughout the 3′-soft clip sequences. To calculate the continuous score, a transition from one residue to the same residue was scored as 0, and a transition from one residue to a different residue scored as 1. The number of A, U, C, and G residues was also counted in the 3′-soft clip sequences of each alignment. The 3′-soft clip sequences with frequencies of U, C, and G all greater or equal to 0.1 were marked as “HIGH_TCG” tails. The 3′-soft clips which were not marked as “HIGH_TCG” and with continuous scores less than or equal to 12 were considered poly(A) tails.

### Poly(A) tail length measurement

To accurately determine the lengths of poly(A) tails, we only quantified the poly(A) tail length from clean CCS reads with at least ten passes. The poly(A) tail length of a transcript was calculated as the length of the sequence, including U, C, or G residues if present. The poly(A) tail length of a gene was represented by the geometric mean of the poly(A) tail length of transcripts with tail length at least 1 nt from the given gene, because poly(A) tail length distribution of a gene follows a lognormal-like distribution^3^.

### Detection of non-A residues in poly(A) tails

To minimize errors introduced by the sequencer, we used clean CCS reads with at least ten passes to identify non-A residues in poly(A) tails. G, C, and U (presented as T in CCS reads) were counted in the poly(A) tail of each CCS read. The percentage of non-A transcripts (CCS reads which contained any non-adenosine residues) of a gene was calculated as the number of CCS reads containing at least one G, C, or U residue divided by the total number of CCS reads derived from the gene. Oligo-U (U≥3) refers to reads which contain at least three consecutive Us, mono-U refers to reads which contain single U but not any two consecutive Us, and U2 refers to reads which contain UU but not any three consecutive Us.

For assigning the positions of U, C, or G residues in poly(A) tails, a given poly(A) tail was first scanned for 3′-end U, C, or G residues, which if present were trimmed from the sequence, then searched for 5′-end U, C, or G residues, which were also trimmed, and finally searched for internal U, C, or G residues.

For calculating the N number for U, C, or G residues, a given poly(A) tail was first searched for the longest consecutive span of U, C, or G residues. The length of sequence before this longest consecutive stretch of U, C, or G residues was considered the N number. If a given poly(A) tail contained multiple stretches of longest consecutive U, C, or G residue, then the N number for this tail could not be determined and thus it was discarded from the N number analysis.

### Analysis of maternal and zygotic genes

Read counts for each gene were summarized using *featureCounts*. The maternal and zygotic genes were defined using the PAIso-seq1 data following a published strategy with minor modifications^48^. In brief, *edgeR* was used for differential expression analysis^49^. The maternal genes were defined by protein coding genes showing 4-fold enrichment in GV oocytes compared to 8C embryos at the transcript level (*p* < 0.05), while the zygotic genes were defined by protein coding genes showing 4-fold enrichment in 8C embryos compared to GV oocytes at the transcript level (*p* < 0.05). Pearson correlations of gene expression between replicates and different samples were calculated using the *cor* function, and the correlation heatmap was generated using pheatmap in *R*.

### Genome and gene annotation

The genome sequence used in this study is from the following links. http://ftp.ebi.ac.uk/pub/databases/gencode/Gencode_human/release_36/GRCh38.primary_assembly.genome.fa.gz

The genome annotation (including the nuclear encoded mRNAs, lncRNAs and mitochondria encoded mRNAs) used in this study is from the following links. http://ftp.ebi.ac.uk/pub/databases/gencode/Gencode_human/release_36/gencode.v36.primary_assembly.annotation.gtf.gz

### Microinjection of siRNA

The set of siRNAs against *BTG4*, *TUT4*, *TUT7*, *TENT4A*, *TENT4B* and non-targeting control siRNA were purchased from Dharmacon. The sequence information of siRNAs is included in Extended Data Table 1. The GV oocytes were microinjected with 5-10 pl siRNA (10 μM) and cultured for the indicated time.

### Data Availability

The ccs data in bam format from PAIso-seq1and 2 experiments will be available at Genome Sequence Archive hosted by National Genomic Data Center. Custom scripts used for data analysis will be available upon request.

## Acknowledgements

We thank Yiwei Zhang for his technical assistance in bioinformatic analysis. This work was supported by the National Key Research and Development Program of China (2020YFA0804000, 2018YFC1004000),the Strategic Priority Research Program of the Chinese Academy of Sciences (XDA24020203, XDA16010113), National Natural Science Foundation of China (31970588, 32170606, 81871168), Natural Science Foundation of Heilongjiang province (YQ2020C003), the China Postdoctoral Science Foundation (2020M670516, 2020T130687), the State Key Laboratory of Molecular Developmental Biology, and the Fundamental Research Funds of Shandong University.

## Author Contributions

Yusheng Liu, Jiaqiang Wang and Falong Lu conceived the project and designed the study. Yusheng Liu, Fanghong Shao, Hu Nie, Jiaqiang Wang, Bing Zhou and Falong Lu analyzed the sequencing data. Keliang Wu, Jingye Zhang, Cheng Li, Zhenzhen Hou and Han Zhao collected human oocytes and embryos, performed drug treatment on human embryos and siRNA mediated knock-down in human oocytes and embryos. Yusheng Liu and Jiaqiang Wang organized all figures. Yusheng Liu, Jiaqiang Wang and Falong Lu supervised the project. Yusheng Liu, Jiaqiang Wang and Falong Lu wrote the manuscript with the input from the other authors.

## Competing Interests statement

The authors declare no competing interests.

---

**Extended Data Fig. 1.**
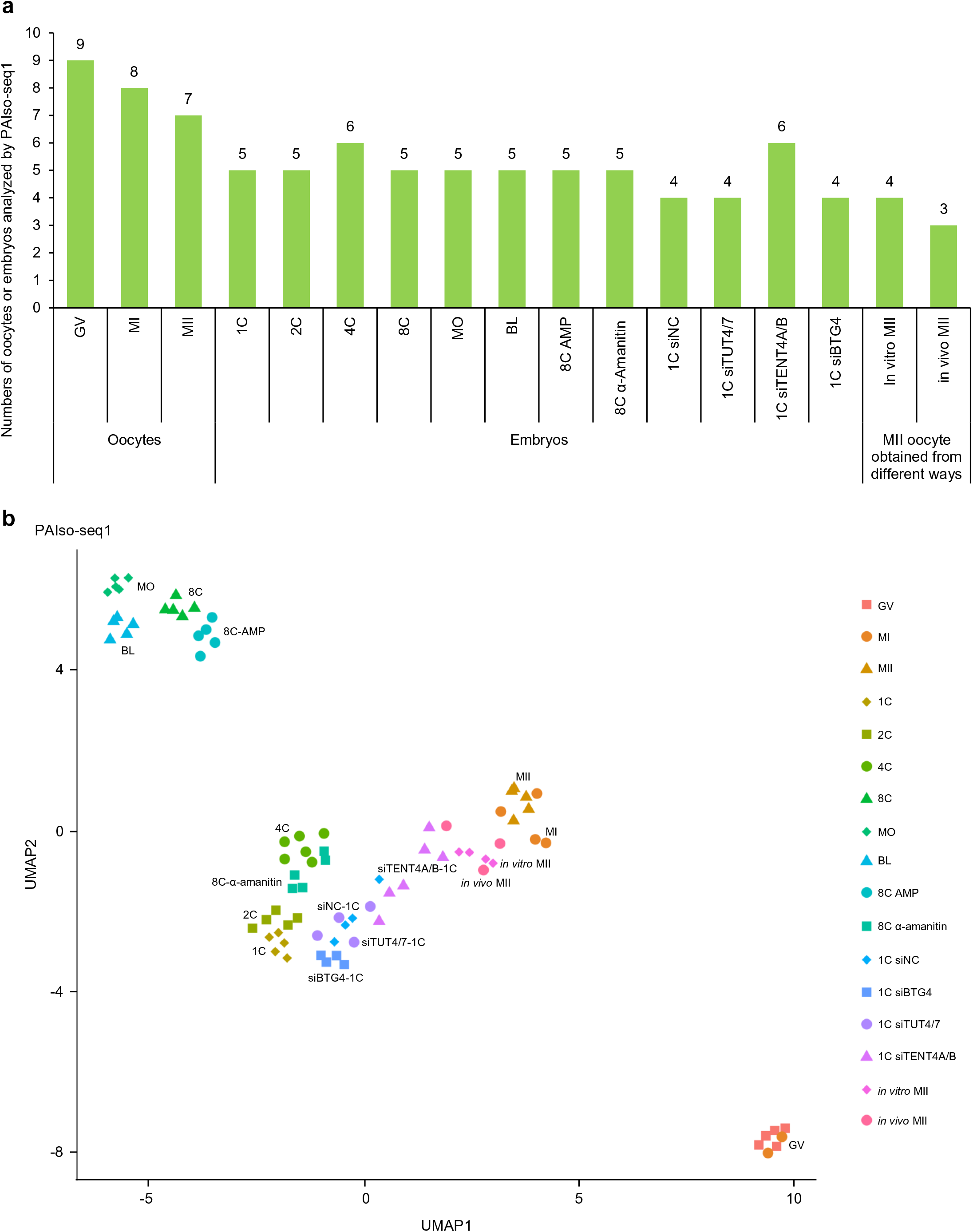
Information on human oocytes and embryos analyzed with PAIso-seq1. **a,** Numbers of single human oocytes and embryos analyzed with PAIso-seq1 for different stages. **b,** The uniform manifold approximation and projection (UMAP) visualization of human oocyte and embryo sample clustering based on expression analyzed with PAIso-seq1.

**Extended Data Fig. 2.**
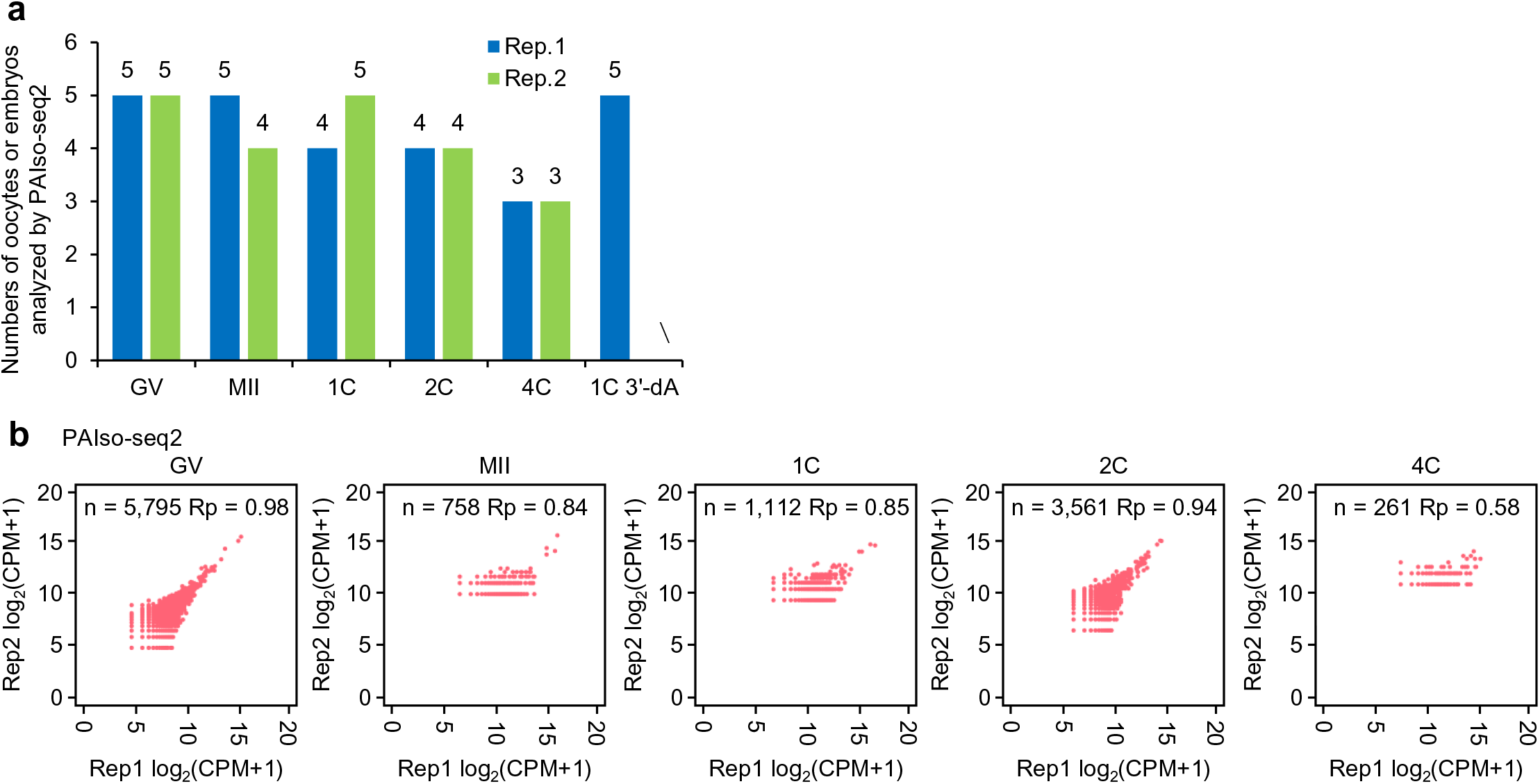
Information on human oocytes and embryos analyzed by PAIso-seq2. **a,** Numbers of human oocytes and embryos analyzed by each PAIso-seq2 replicate for different stages. **b,** Scatter plots showing the Pearson correlation of gene expression between two replicates for human oocytes and embryos at different stages measured with PAIso-seq2. Each dot represents one gene. Pearson’s correlation coefficient (Rp) and number of genes included in the analysis are shown at the top.

**Extended Data Fig. 3.**
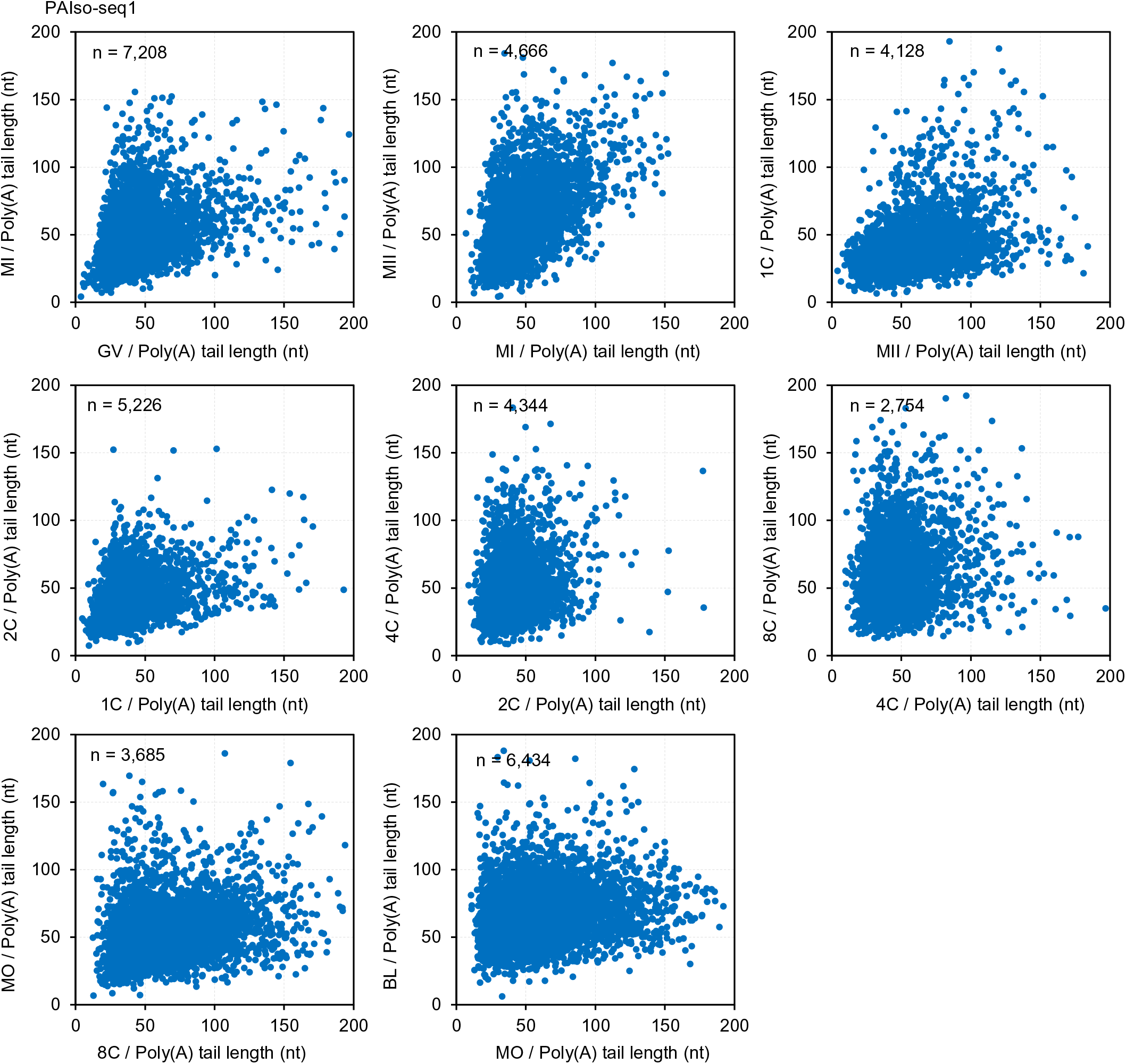
The change in poly(A) tail length for each gene between two neighboring developmental stages measured with PAIso-seq1. Scatter plots of poly(A) tail length of human samples at neighboring developmental stages measured with PAIso-seq1. Each dot represents one gene. The poly(A) tail length for each gene is the geometric mean length of all the transcript with poly(A) tail at least 1 nt in length for the given gene. For each of the graphs, genes with at least 10 reads in both of the samples were included in the analysis. The number of genes included in the analyses is included at the top left of the graphs.

**Extended Data Fig. 4.**
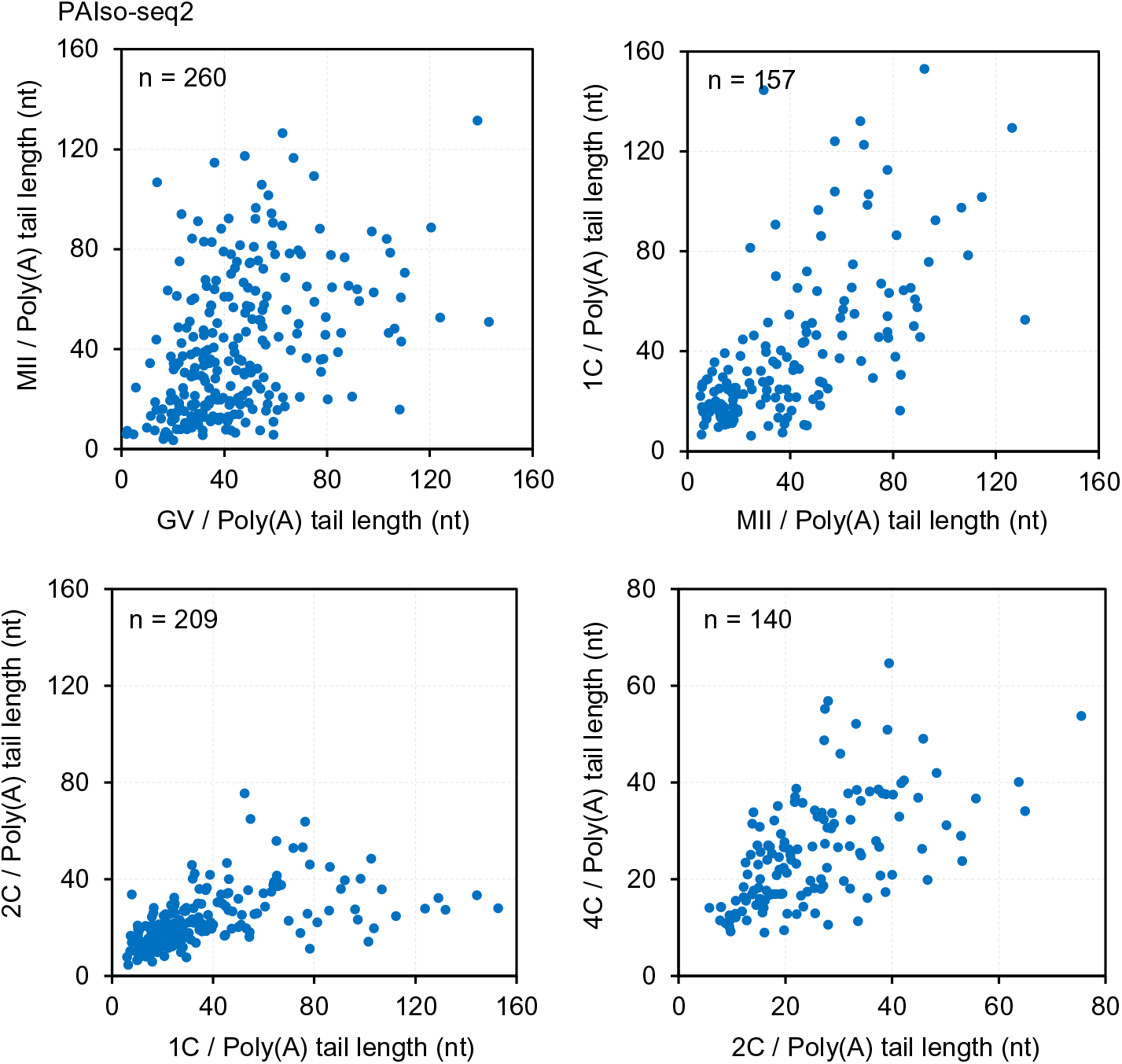
The change in poly(A) tail length for each gene between two neighboring developmental stages measured with PAIso-seq2. Scatter plots of poly(A) tail length of human samples at neighboring developmental stages measured with PAIso-seq2. Each dot represents one gene. The poly(A) tail length for each gene is the geometric mean length of all the transcript with poly(A) tail at least 1 nt in length for the given gene. For each of the graphs, genes with at least 10 reads in both of the samples were included in the analysis. The number of genes included in the analyses is included at the top left of the graphs.

**Extended Data Fig. 5.**
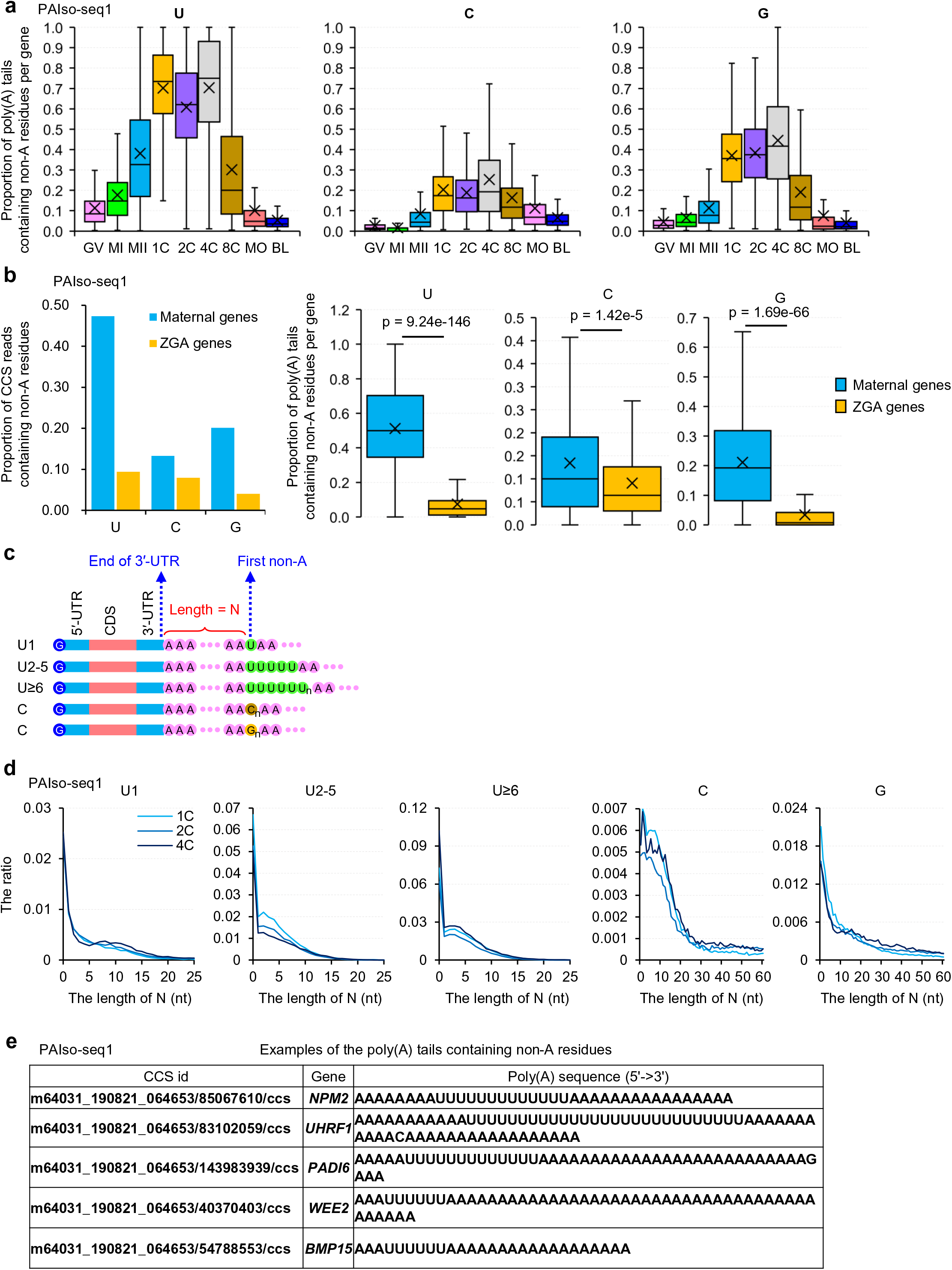
Characterization of non-A residues during the human OET measured with PAIso-seq1. **a,** Box plot for the proportion of reads containing U, C, or G residues of individual genes in samples at different stages measured with PAIso-seq1. Genes (n = 1,664) with at least 10 poly(A) tail containing reads (tail length ≥ 1 nt) were included in the analysis. **b,** Overall proportion of transcripts containing U, C or G residues for combined transcripts from maternal genes (n = 3,339) or zygotic genes (n = 1,502) in 8C embryos measured with PAIso-seq1 (left). Box plot of the proportion of reads containing U, C, or G residues for each of the maternal genes (n = 118) or zygotic genes (n = 640) in 8C embryos measured with PAIso-seq1 (right). Genes with at least 20 poly(A) tail containing reads (tail length ≥ 1 nt) were included in the analysis. The *p* values were calculated by Student’s *t* tests. **c,** Diagram depicting mRNA with 5′-end or internal non-A residues. N represents the length of residues between the end of 3′ UTR and the first base of the longest consecutive U, C, or G residue sequence in a poly(A) tail. **d,** Histogram of the length of N and the ratio of U1, U2-5, U≥6, C, and G residues in human 1C, 2C, and 4C embryos measured with PAIso-seq1. Histograms (bin size = 1 nt) are normalized to the total number of transcripts with poly(A) tail at least 1 nt in length. **e,** Examples of poly(A) tails with internal consecutive U residues in human zygotes measured with PAIso-seq1. For all box plots, the “×” indicates the mean value, black horizontal bars show the median value, and the top and bottom of the box represent the values of the 25^th^ and 75^th^ percentiles, respectively.

**Extended Data Fig. 6.**
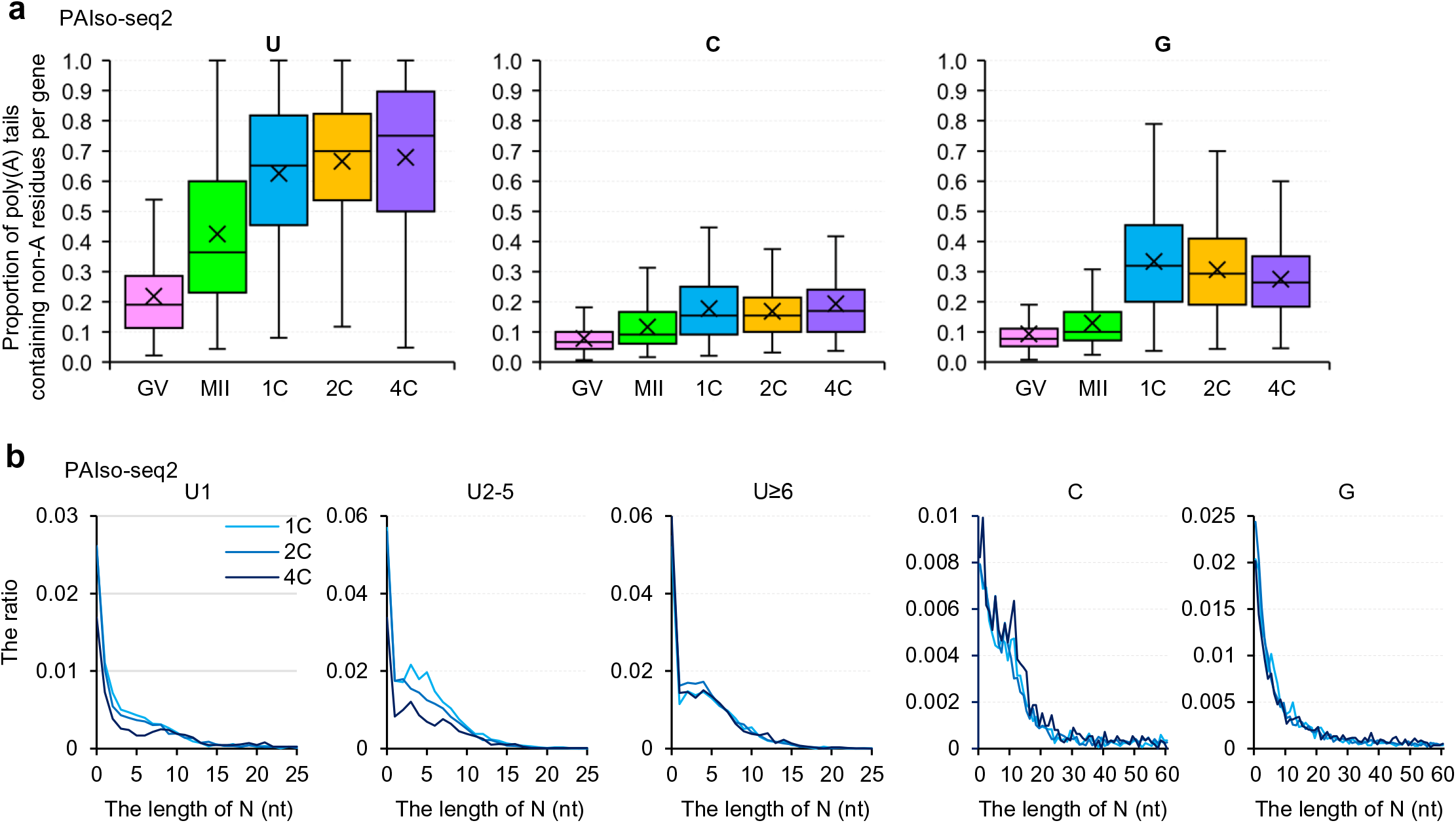
Characterization of non-A residues during the human OET measured with PAIso-seq2. **a,** Box plot of the proportion of reads containing U, C, or G residues of individual genes in samples at different stages measured with PAIso-seq2. Genes (n = 100) with at least 10 poly(A) tail containing reads (tail length ≥ 1) were included in the analysis. The “×” indicates the mean value, black horizontal bars show the median value, and the top and bottom of the box represent the values of the 25^th^ and 75^th^ percentiles, respectively. **b,** Histogram of the length of N (see illustration in Extended Data Fig. 5c) and the ratio of U1, U2-5, U≥6, C, and G residues in human 1C, 2C, and 4C embryos measured with PAIso- seq2. Histograms (bin size = 1 nt) are normalized to the total number of transcripts with poly(A) tail at least 1 nt in length. Histograms (bin size = 1 nt) are normalized to the total number of transcripts with poly(A) tail at least 1 nt in length.

**Extended Data Fig. 7.**
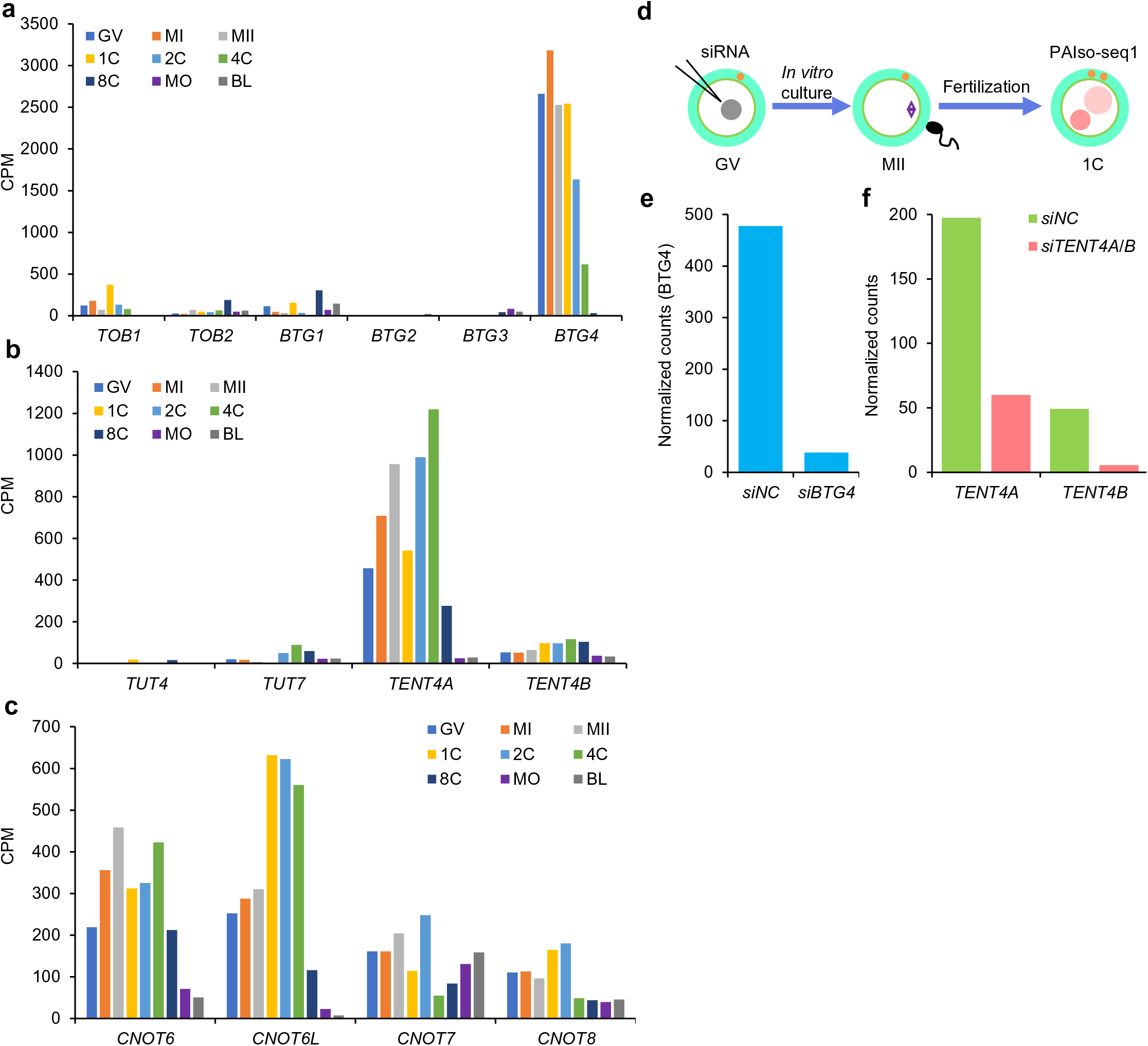
Knockdown of candidate factors involved in poly(A) tail regulation. **a,** The expression pattern of *TOB1*, *TOB2*, *BTG1*, *BTG2*, *BTG3*, and *BTG4* genes in human different stage oocytes and embryos revealed with PAIso-seq1 data. **b,** The expression pattern of *TUT4*, *TUT7*, *TENT4A*, and *TENT4B* genes in human different stage oocytes and embryos revealed with PAIso-seq1 data. **c,** The expression pattern of *CNOT6*, *CNOT6L*, *CNOT7*, and *CNOT8* family genes in human different stage oocytes and embryos revealed with PAIso-seq1 data. **d,** Illustration of siRNA microinjection followed by *in vitro* maturation and *in vitro* fertilization. **e,** Normalized read counts of *BTG4* in *siNC* and *siBTG4* human zygotes measured with PAIso-seq1. **f,** Normalized read counts of *TENT4A* and *TENT4B* in *siNC* and *siTENT4A/B* human zygotes measured with PAIso-seq1. The read counts were normalized by counts of reads mapped to protein-coding genes in the mitochondria genome if normalization was indicated.

**Extended Data Fig. 8.**
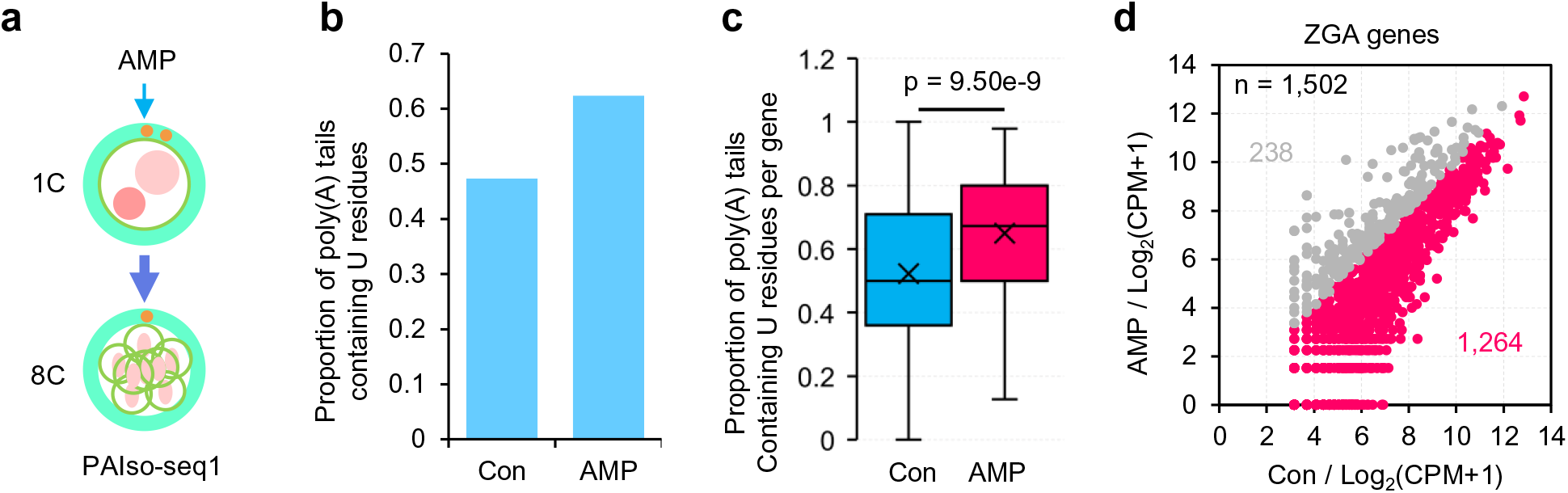
AMP treatment in human 8C embryos. **a,** Illustration of the embryo AMP treatment experiments. **b,** Proportion of combined transcripts from maternal genes (n = 3,339) containing U residues in human 8C embryos with or without AMP treatment measured with PAIso-seq1. Transcripts with poly(A) tail at least 1 nt in length were included in the analysis. **c,** Box plot of the proportion of reads containing U residues for individual maternal genes (n = 3,339) in human 8C embryos with or without AMP treatment measured with PAIso- seq1. Transcripts with poly(A) tail at least 1 nt in length were included in the analysis. **d,** Scatter plot of the expression level of ZGA genes in human 8C embryos with and without AMP treatment measured with PAIso-seq1. Each dot represents one gene. Genes with at least 1 read in one of the samples were included in the analysis. The number of genes included in the analyses is included at the top left of the graphs. All *p* values were calculated by Student’s *t* tests. For all box plots, the “×” indicates the mean value, black horizontal bars show the median value, and the top and bottom of the box represent the values of the 25^th^ and 75^th^ percentiles, respectively.

## Notes

### Competing Interest Statement

The authors have declared no competing interest.

